# Identification of a novel interspecific hybrid yeast from a metagenomic spontaneously inoculated beer sample using Hi-C

**DOI:** 10.1101/150722

**Authors:** Caiti Smukowski Heil, Joshua N. Burton, Ivan Liachko, Anne Friedrich, Noah A. Hanson, Cody L. Morris, Joseph Schacherer, Jay Shendure, James H. Thomas, Maitreya J. Dunham

## Abstract

Interspecific hybridization is a common mechanism enabling genetic diversification and adaptation; however, the detection of hybrid species has been quite difficult. The identification of microbial hybrids is made even more complicated, as most environmental microbes are resistant to culturing and must be studied in their native mixed communities. We have previously adapted the chromosome conformation capture method Hi-C to the assembly of genomes from mixed populations. Here, we show the method’s application in assembling genomes directly from an uncultured, mixed population from a spontaneously inoculated beer sample. Our assembly method has enabled us to de-convolute 4 bacterial and 4 yeast genomes from this sample, including a putative yeast hybrid. Downstream isolation and analysis of this hybrid confirmed its genome to consist of *Pichia membranifaciens* and that of another related, but undescribed yeast. Our work shows that Hi-C-based metagenomic methods can overcome the limitation of traditional sequencing methods in studying complex mixtures of genomes.

## Introduction

Once thought to be a rare phenomenon, hybrids are increasingly recognized as common in natural and industrial environments (Abbott et al. 2013). Hybridization can act as an instantaneous, large-scale mutational event, introducing abundant genetic variation and potentially producing changes in gene expression, gene content, and karyotype. New combinations of alleles formed during the initial hybridization event, or transgressive segregation from subsequent recombination, may allow the hybrid to explore greater phenotypic space than either of its parents, and thus to colonize new, uninhabited niches. There is abundant evidence suggesting that because hybrids can explore unoccupied ecological niches, they may be particularly adept at invading novel habitats and even become invasive pests (Bullini 1994; Mallet 2007; Muhlfeld et al. 2014; Pryszcz et al. 2014). In line with these findings, extensive hybridization is often associated with human-disturbed habitats (Dowling and Secor 1997; Vallejo-Marin and Hiscock 2016), and climate change is predicted to increase the number of hybridization events (Kelly et al. 2010; Hoffmann and Sgro 2011).

Similarly, hybrids are often found in commercial settings where stress tolerance or adaptation to particular environmental conditions is important. Most famously, the lager brewing yeast *Saccharomyces pastorianus* is a hybrid between the commensal baking and brewing yeast, *Saccharomyces cerevisiae*, and the cryotolerant species *Saccharomyces eubayanus. S. cerevisiae* and *S. eubayanus* diverged approximately 20 million years ago and are 20% divergent at the coding level (Tamai et al. 1998; Yamagishi and Ogata 1999; Dunn and Sherlock 2008; Nakao et al. 2009; Baker et al. 2015; Gibson and Liti 2015). After what is presumed to be an allopolyploid event, *S. pastorianus* has undergone massive loss of chromosomes and chromosomal segments. While *S. pastorianus* genomic content remains fairly balanced in representation of both parental genomes, many hybrids are observed to have imbalanced retention and loss patterns. Hybrids amongst other members of the *Saccharomyces* clade have been discovered in wine fermentation and brewing operations (Gonzalez et al. 2006; Gonzalez et al. 2008; Bellon et al. 2015; Wolfe 2015; Magalhaes et al. 2017), in addition to a number of other fungal species hybrids outside of this well-studied clade. For example, a large proportion of strains of the spoilage yeast *Dekkera* (Brettanomyces) *bruxellensis*, which itself is used in brewing, originated through interspecific hybridization (Hellborg and Piskur 2009; Borneman et al. 2014), and these hybrids have a selective advantage over non-hybrids in wineries (Borneman et al. 2014).

It has been suggested that the previously unrecognized ubiquity of hybrid species is at least in part due to issues of detection and identification (Dowling and Secor 1997), which may be particularly true in microbial taxa (Albertin and Marullo 2012). One of the issues complicating detection of microbial hybrids is the difficulty of dissecting microbial communities in general, which can be made up of many different species, most of which are unculturable (Lane et al. 1985; Pace et al. 1986; Handelsman 2004; Marcy et al. 2007). With the advancement of next generation sequencing, the field of metagenomics has moved from relying on 16S rRNA gene fingerprinting methods (Lane et al. 1985; Pace et al. 1986; Rajendhran and Gunasekaran 2011) to various applications of shotgun sequencing (Venter et al. 2004; Human Microbiome Project 2012; Iverson et al. 2012; Howe et al. 2014), single-cell genome sequencing (Marcy et al. 2007; Stepanauskas 2012; Kamke et al. 2013), and associated computational methods (Saeed et al. 2012; Carr et al. 2013; Hug et al. 2013; Wood and Salzberg 2014).

Previous work from our groups and others demonstrated the use of contact probability maps generated through chromosome conformation capture methods (Lieberman-Aiden et al. 2009; Dekker et al. 2013) to assemble complex genomes (Burton et al. 2013; Kaplan and Dekker 2013), and in particular, to deconvolute complex metagenomic samples (Beitel et al. 2014; Burton et al. 2014; Marbouty et al. 2014). The basis of this idea takes advantage of the Hi-C protocol, in which crosslinking of proximal chromatin segments occurs prior to cell lysis, thereby ensuring that each Hi-C interaction involves a pair of reads generated from within the same cell. In a diverse microbial population, such data can help delineate which genomic sequences originated from individual species. Burton et al. demonstrated that the method, MetaPhase, can reliably deconvolute and assemble genomes from up to 18 eukaryotic and prokaryotic species from synthetic mixtures in the lab. Importantly, this method should also allow hybrids to be identified in a mixed sample, since crosslinks would be made between the different genomes in contrast to the solely within-genome linkages in a simple mix of the parental species.

Here, we extend this work to explore the microbial community of a spontaneously inoculated beer sample. In the tradition of Belgian lambic, boiled wort is left open to the elements and is fermented solely from autochthonous yeast and bacteria (Vanoevelen et al. 1977; Verachtert and Iserentant 1995; Keersmaecker 1996). Past studies of lambic style fermentation have relied heavily on culturing various species over the course of fermentation; more recently, genetic tools such as terminal restriction fragment length polymorphism, barcoded amplicon sequencing, and quantitative PCR have shed light on the microbial progression, but still are limited by deconvolution issues (Bokulich et al. 2012; Spitaels et al. 2014c). Our method deconvolutes the microbial genomes in the beer sample without culturing or a priori knowledge of species or strain, revealing eight organisms, including novel strains and species of bacteria and yeast, and a novel yeast hybrid. The hybrid appears to be a recent hybridization event between *Pichia membranifaciens* and a previously undescribed species approximately 10-20 million years divergent. We characterize the genome and fermenting capabilities of this novel hybrid, and describe the other species identified. This is one of the first demonstrations of computational metagenomic deconvolution of a non-laboratory sample (see also Marbouty et al. 2017 and Marbouty et al. 2014), and the first method with the ability to detect hybrids in a heterogeneous population.

## Materials & Methods

### Sample collection

We obtained 20 mL of actively growing culture sampled from the top of a wine barrel containing the spontaneously inoculated beer “Old Warehouse,” produced by Epic Ales in Seattle, WA on May 8, 2014.

### Shotgun, Hi-C libraries

Approximately 5 mL of the sample was pelleted and total DNA was isolated using a standard phenol/chloroform glass bead extraction. Shotgun libraries were prepared using the Nextera Kit (Illumina). Hi-C libraries were prepared as described (Burton et al. 2014). Sequencing was performed on the NextSeq 500 Illumina platform.

### De novo assembly, deconvolution, and individual species assembly

The draft metagenome assembly was created using the IDBA-UD assembler (Peng et al. 2012) with the following parameters: ‘–pre_correction –mink 20 –maxk 60 –step 10’. Hi-C reads were aligned to the draft assembly using BWA (Li and Durbin 2009), following the strategy of Burton et al. (2014). Clustering of contigs into individual clusters was done using MetaPhase (Burton et al. 2014); https://github.com/shendurelab/MetaPhase). Independently of MetaPhase, in order to determine species identity, contigs were mapped to the BLAST sequence database (https://blast.ncbi.nlm.nih.gov/Blast.cgi; July 2014 database), using blastn with the following parameters:-perc_identity 95-evalue 1e-30-word_size 50. Percentage of each cluster assembly aligning to the reference was estimated using AssemblyEvaluator (https://github.com/snayfach/AssemblyEvaluator).

### Hybrid assembly and analysis

To confirm the putative hybrid from the metagenome assembly, the hybrid was isolated from a single colony. DNA was extracted and prepared with a Nextera kit, as above. Reads were mapped to the metagenome assembly using BWA (Li and Durbin 2009), and once confirmed as a hybrid, a new draft assembly was created using IDBA-UD with parameters as above (Peng et al. 2012). To split the assembly into species-specific sub-genomes, contigs from this new assembly were compared against the *P. membranifaciens* genome v2.0 (Riley et al. 2016) using blastn with an e-value of 1E-12 (Altschul et al. 1990). All contigs whose single best blastn match to the *P. membranifaciens* genome is >= 97% identical were classified as sub-genome A, whereas all contigs whose single best blastn match to the *P. membranifaciens* genome is >= 77% and < 92% identical were classified as sub-genome B. Contigs of high divergence and contigs with identity between 92-97% were unable to be parsed into sub-genomes. Augustus v3.2.1 was used to create gene predictions for both sub-genomes (Stanke et al. 2004; Stanke and Morgenstern 2005). Gene predictions between sub-genomes were compared using blastn and blastp using the best hit. BLAST was also used to assess coverage differences between the subgenomes. A rough approximation of divergence between the sub-genomes was estimated by constructing neighbor-joining trees for *P. membranifaciens* predicted genes (g1201, g2530) using ClustalOmega (Sievers et al. 2011) and PHYLIP (Felsenstein 2005), and sequence from each sub-genome, as compared with results using *S. cerevisiae, S. paradoxus, S. kudriavzevii,* and *S. uvarum*.

### *Neighbor-joining tree building for* S. cerevisiae *strain phylogeny*

The biallelic segregating sites detected among a set of previously sequenced *S. cerevisiae* isolates (Strope et al. 2015; Gallone et al. 2016) and the OW sequence were used to elucidate the phylogenetic relationships among them. Based on these sites, a distance matrix was first computed and submitted to bionj algorithm (R packages ape and SNPrelate) to construct a neighbor-joining tree.

### Lab-appropriate wort

Our protocol was designed to make 1.75 liters of a lab-appropriate American style amber beer wort. 320 grams of an amber liquid malt extract are used to meet the Brewers Association's Beer Standard Guidelines for an American style amber/red ale with an original gravity of 11.9 - 14.3 °Plato. Our wort was within this range at 13.77 °Plato. The sugar composition was: 44% maltose, 11% maltotriose, 13% glucose, 3% sucrose, 4% fructose, and 25% unfermentable dextrins. To prepare media, 320 grams of liquid malt extract was added to 1.5 liters of 77 °C water using a heat block with a running stir bar. Once in solution, the heat was adjusted such that the wort was allowed to achieve a rolling boil without overflowing for 1 hour. The media was then repeatedly poured through paper filters until achieving a uniform, cloudless consistency. Distilled water was added to bring the solution to a total of 1.75 liters before being covered. The media was vacuum filtered through Nalgene 500mL Rapid-Flow Bottle Top Filters with a pore size of 0.45μm, and into a sterile 2L bottle. The bottle was stored at room temperature and away from light exposure. Wort was inoculated with a single colony suspended in 2 mL of water after the temperature reached 22 °C. The wort was aerated via swirling at the point of inoculation, after which no shaker was used and no air was pumped in. It was kept at room temperature for the duration of 5 weeks. After 5 weeks, ethanol volume, pH, and sugar content were analyzed (White Labs, San Diego, CA). No single sugar fermentation studies were conducted at this time.

### Data availability

Reads and assemblies for the metagenome are deposited at NCBI under the Project ID PRJNA390460.

## Results

### Hi-C/MetaPhase on a craft beer sample

We obtained a 20mL sample of active culture growing in wort that had been inoculated in open air and serially pitched in a used wine barrel for several years by Epic Ales in Seattle, Washington. We performed shotgun sequencing and Hi-C on this sample and applied the approach utilized by Burton et al. (2014) to assemble and deconvolute the metagenomic sample. Briefly, we sequenced two shotgun libraries, one with 3.61 M read pairs of 35 bp each, and one with 27.5 M read pairs of 100 bp each, and two Hi-C libraries, one using the HindIII restriction site and one using the NcoI restriction site, with 13.6 M and 14.5 M read pairs respectively. We converted the shotgun reads into a *de novo* assembly using IDBA-UD (Peng et al. 2012), yielding 32,549 contigs with an N50 of 5,285 bp at a minimum contig length of 200 bp and a total size of 67,496,013 bp. We mapped the Hi-C reads to the assembly using BWA (Li and Durbin 2009), and applied the MetaPhase software to deconvolute draft contigs into a progressively decreasing number of clusters while measuring intra-cluster enrichment signal as described in Burton et al. (2014). This signal reached a maximum at *N=8* clusters (**Figure 1B**), suggesting the presence of 8 discernable species-level genomes in this assembly (**Figure 1A, Table 1**). 11,189 contigs are included in these eight clusters, representing 73.3% of the initial sequence assembly (49,498,624 bp); the remaining contigs are small (N50 of 1349 bp) and were unable to be confidently clustered. Seven of the eight clusters matched to a unique species of either fungal or bacterial origin (**Figure 1C**), and in each of these clusters, the quantity of sequence served to indicate the completeness of coverage of the species’ genome (**Table 1**). In the eighth cluster, no significant quantity of sequence closely matched a known reference genome; however, several sequences in this cluster matched individual genes from *Pediococcus damnosus*, a known spoilage bacterium for which no complete reference genome existed in the BLAST database at the time of analysis (but see (Snauwaert et al. 2015)). This cluster is now confirmed to be *P. damnosus*.

**Figure 1:**
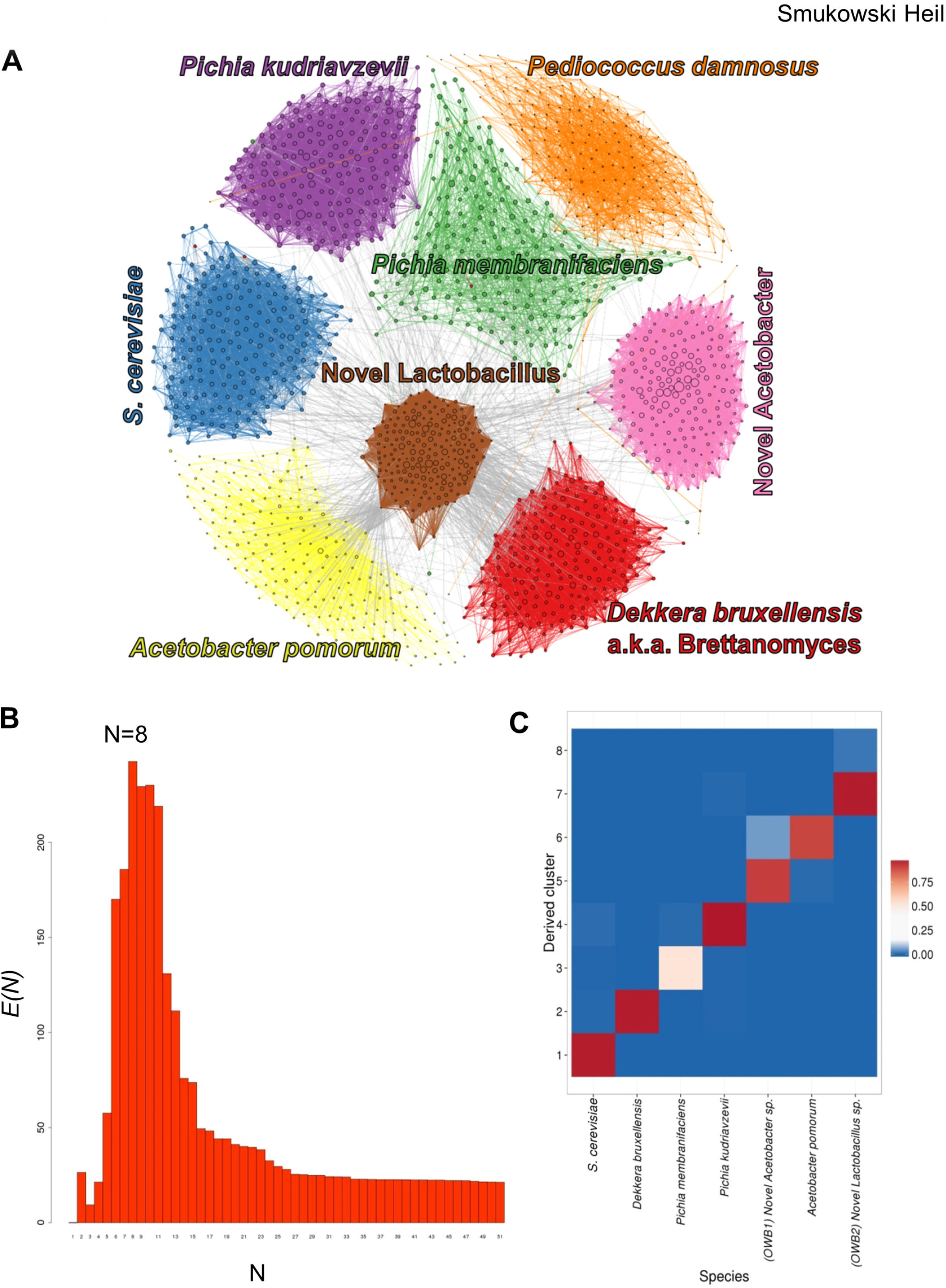
MetaPhase deconvolutes species from a spontaneously inoculated beer sample. **A.** Clustering of the Old Warehouse assembly. Each contig is shown as a dot, with size indicating contig length, colored by species. Edge widths represent the densities of Hi-C links between the contigs shown. **B.** Determining the approximate number of species in the metagenome. We ran the hierarchical agglomerative clustering algorithm on the draft assembly. In this algorithm, the number of clusters starts high and gradually decreases as clusters are merged together; to generate this data, we continued clustering all the way down to N = 1 cluster. Shown is the metric *E*, or intra cluster link enrichment, at each value of *N*. It is assumed that the maximum value of *E(N)* occurs when *N* is roughly equal to the true number of distinct species present in the draft assembly. For this assembly, this occurs at *N* = 8 clusters. **C.** Validation. This heatmap indicates what fraction of the sequence in each MetaPhase cluster maps uniquely to each of the reference genomes of the predicted species. Note that not all sequence is expected to map uniquely to one species. x-axis: the 7 species. y-axis: the MetaPhase clusters. Note, this excludes the *P. damnosus-like* cluster because there was no public *P. damnosus* reference genome available.

**Table 1:**
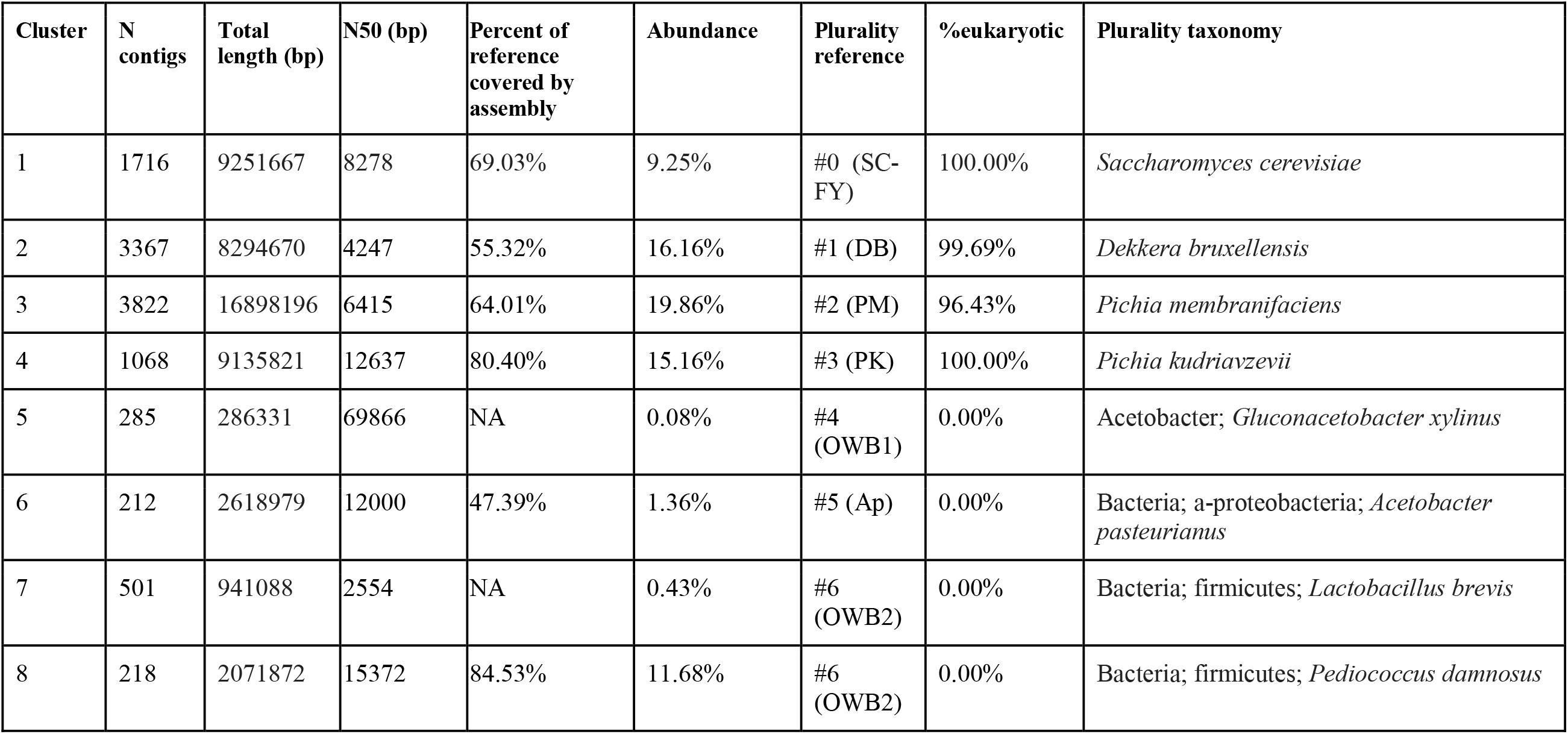
Species identified in spontaneously inoculated beer.

| Cluster | N contigs | Total length (bp) | N50 (bp) | Percent of reference covered by assembly | Abundance | Plurality reference | %eukaryotic | Plurality taxonomy |
| --- | --- | --- | --- | --- | --- | --- | --- | --- |
| 1 | 1716 | 9251667 | 8278 | 69.03% | 9.25% | #0 (SC-FY) | 100.00% | *Saccharomyces cerevisiae* |
| 2 | 3367 | 8294670 | 4247 | 55.32% | 16.16% | #1 (DB) | 99.69% | *Dekkera bruxellensis* |
| 3 | 3822 | 16898196 | 6415 | 64.01% | 19.86% | #2 (PM) | 96.43% | *Pichia membranifaciens* |
| 4 | 1068 | 9135821 | 12637 | 80.40% | 15.16% | #3 (PK) | 100.00% | *Pichia kudriavzevii* |
| 5 | 285 | 286331 | 69866 | NA | 0.08% | #4 (OWB1) | 0.00% | Acetobacter; *Gluconacetobacter xylinus* |
| 6 | 212 | 2618979 | 12000 | 47.39% | 1.36% | #5 (Ap) | 0.00% | Bacteria; a-proteobacteria; *Acetobacter pasteurianus* |
| 7 | 501 | 941088 | 2554 | NA | 0.43% | #6 (OWB2) | 0.00% | Bacteria; firmicutes; *Lactobacillus brevis* |
| 8 | 218 | 2071872 | 15372 | 84.53% | 11.68% | #6 (OWB2) | 0.00% | Bacteria; firmicutes; *Pediococcus damnosus* |

### Identification and characterization of a hybrid yeast

One of the more abundant organisms in the sample had sequencing reads from the yeast *Pichia membranifaciens*, a common food spoilage species (Kurtzman et al. 2011; Riley et al. 2016). *P. membranifaciens* has been identified in spontaneously inoculated beer (Spitaels et al. 2014c), wine (Rankine 1966; Stefanini et al. 2016a), and other beverages (Lachance 1995; Silva et al. 2009). The assembly generated here has a genome size roughly twice that of *P. membranifaciens*, and upon further inspection, BLAST results were consistent with two separate genomes, one belonging to *P. membranifaciens*, and the other to an unknown, related species. We hypothesized that this could result either from a technical error in MetaPhase collapsing two species into one cluster, or alternatively, a hybridization event. To test this hypothesis, isolates of the putative hybrid were grown and sequenced (**Figure 2A**). Sequencing reads were mapped back to the MetaPhase assembly, and mapped exclusively to the cluster identified as the hybrid or to unclustered reads (which were unable to be assembled), substantiating cluster 3 as a true hybrid. This is further supported by ploidy analysis confirming a genome size of approximately 24 Mb (**Figure 2B**).

**Figure 2:**
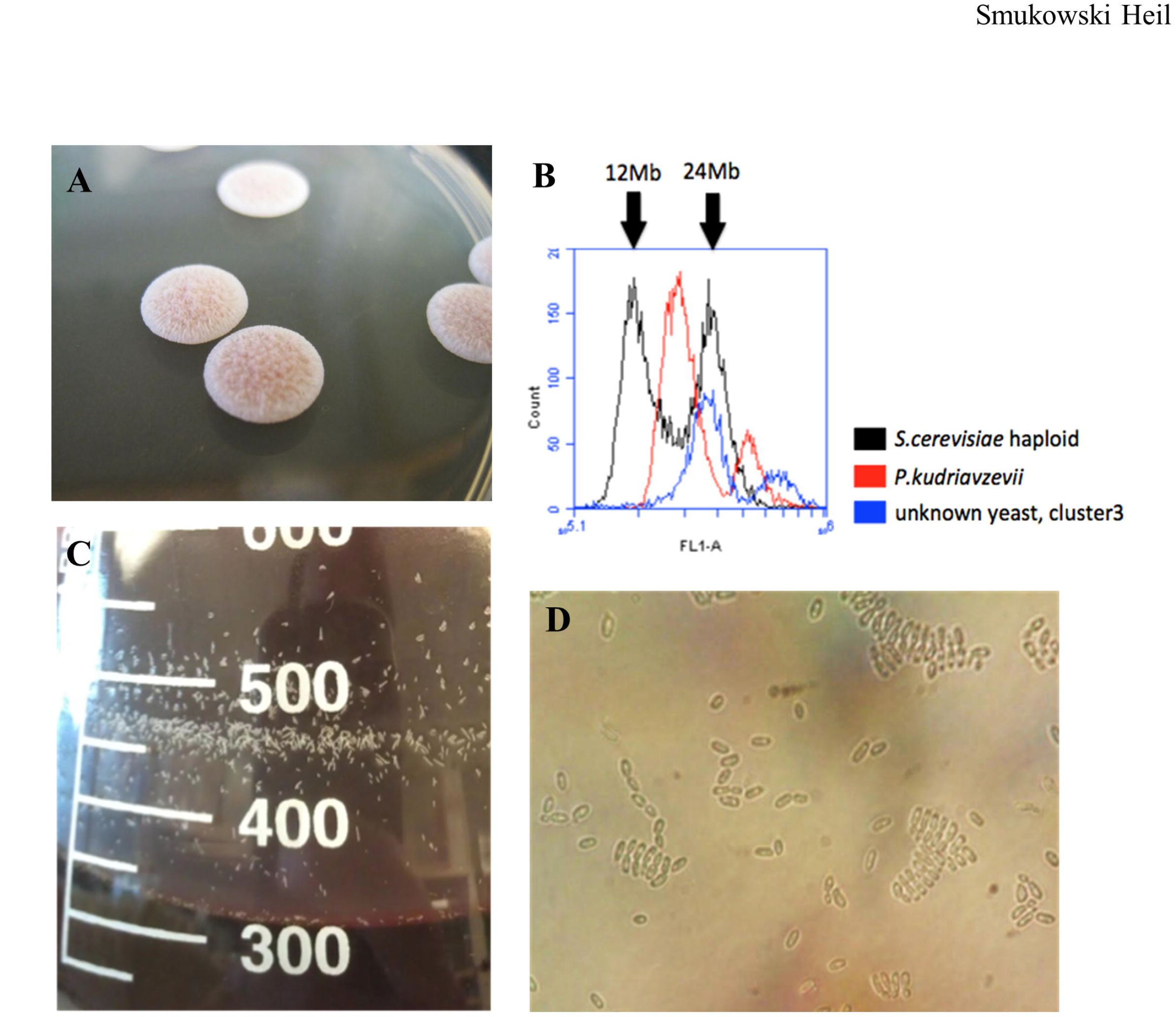
A novel interspecific *Pichia* hybrid. **A.** Isolated colonies of the Pichia hybrid show distinct colony morphology. **B.** Flow cytometry indicates the hybrid genome (blue) size is approximately 24 Mb, similar to G2 haploid *S. cerevisiae*. **C.** Growth in wort shows aggregates on the flask side. **D.** Growth in wort shows palisades pattern.

Contigs from the assembly were parsed into two genomes using percent identity compared to *P. membranifaciens* (Riley et al. 2016), with contig identity above 97% classified as sub-genome A *(P. membranifaciens)*, and contig identity between 77-92% classified as subgenome B (unknown species). Sub-genome A has 5792 predicted genes and sub-genome B has 5742 predicted genes, similar to the 6604 ORFs in *S. cerevisiae* (of which 5159 have been verified). Of 4780 unique protein matches between the two sub-genomes, 92% of proteins share 70% amino acid similarity or higher (4410 proteins), and of these, 56% have greater than 90% amino acid identity. Phylogenetic analysis suggests approximately 10-20 million years divergence between the hybrid parentals, which appears consistent with the amino acid divergence. However, we note that this could be an underestimate since we only considered genes with matches between sub-genomes.

Many hybrids experience gene loss, copy number changes, and aneuploidy following a hybridization event (Abbott et al. 2013). In a crude sense, the amount of loss can date a hybridization event, with more genomic homogenization indicating a more ancient hybridization event. To estimate the “completeness” of the hybrid genome, hybrid contigs were compared to *P. membranifaciens* to obtain coverage estimates. About 2.5% of the *P. membranifaciens* genome has no coverage from the hybrid, mostly in telomeric regions. Of the rest of the genome, 88.8% is covered exactly twice, and 89.6% is covered two or more times. Most of the regions with greater than 2X coverage are likely artifacts of the blast method and overlapping assembly contigs, although we have evidence for potential copy number variants in gene regions in both sub-genomes.

Sub-genome B contig 60_487 is duplicated relative to the rest of the genome and contains three predicted genes: g1333 (methionine permease, *MUP1)*, g1334 (hypothetical protein in *P. membranifaciens*, similarity to methionine permease in *P. kudriavzevii)*, and g1335 (hypothetical protein in *P. membranifaciens*, similar to allantoate permease in *P. kudriavzeviî)*. All three of these genes are present in single copy in sub-genome A. Sub-genome A contig 60_428 is also duplicated and contains a single predicted gene, g1144, an oligopeptide transporter *(OPT2)*. It is not present in sub-genome B, either representing a loss of heterozygosity event, or incomplete assembly. Breakpoints of both duplication events are not identifiable as the copy number increase spans the whole contig, suggesting that the breakpoints are in unassembled regions which may contain additional genes of interest.

Approximately 10.4% of the *P. membranifaciens* genome is covered only once, but the large majority of that is in short fragments distributed across the genome (average length: 202.19 bp, median length: 106 bp), most likely an artifact of incomplete hybrid assembly or unusually divergent regions of subgenome B. Still, there are 120 segments of 1 kb or more that are covered once, representing 2.7% of the genome (total length of 317 kb) and might represent missing genes in one or the other sub-genomes. Overall, these observations are consistent with a recent hybridization event.

### Pichia hybrid does not play a significant role in fermentation

To characterize the fermenting capabilities of the *Pichia* hybrid, the isolate was used to inoculate a wort media at room temperature for five weeks. During this time, aggregates accumulated and could be seen rising to the wort’s surface and attaching to walls of the fermentation vessel (**Figure 2C**). When viewed under a microscope, cells appeared to bend at the points of cell division, resulting in a palisade arrangement with stacking patterns (**Figure 2D**). No pellicles formed on the surface of the wort, in contrast to both *P. membranifaciens* and *P. kudriavzevii,* indicating that the hybrid may have lost this phenotype (Matthew Bochman, pers. comm.).

Filtrates of the starting wort media and the culture after five weeks were analyzed. The pH of the starting wort and the finished product reflect standard brewing measurements at 4.57 and 4.53 pH, respectively. These pH values are consistent with the avoidance of flavor and contamination problems (for example, a higher pH would signal food safety issues), indicating that the *Pichia* hybrid isn’t typical of a spoilage species in this respect.

Extract analyses were completed to determine the amount of non-fermentable carbohydrates remaining in the product after culturing. According to the Real Extract test, the original wort was valued at 13.75 °Plato and after 5 weeks at 13.61 °Plato. The Apparent Extract/Final Gravity in an American style amber/red ale should be 2.5 - 4.6 °Plato (Papazian 2017). The alcohol by weight in this style of beer should be 3.50% - 4.80%, whereas the *Pichia* hybrid product was 0.02%. The final calorie content per a 12 oz. serving of beer typically drops to approximately 160 kcal, whereas the hybrid fermentation product was valued at 202.68 kcal.

In addition to this, percentages based on the entire contents of the media showed that over five weeks, glucose levels dropped from an initial measure of 1.62% to a final measure of 1.22%. The hybrid may incrementally breakdown maltotriose and fructose, dropping from 1.46% to 1.05% and 0.57% to 0.22% respectively, but did not appear to be able to reduce the levels of maltose. However, this test would not distinguish sugars created as metabolic byproducts from those present initially. Single sugar fermentation studies could be done in the future to better define the metabolic activities of the hybrid. These results do indicate that the *Pichia* hybrid did not significantly metabolize much of the available carbohydrates into alcohol within this wort environment, and may suggest that it’s dependent on byproducts or enzymes from other organisms in a mixed population. These results may also reflect a lack of aeration, as we did not employ any aeration methods during the five week fermentation, and a recent study noted *P. kudriavzevii* is not able to grow in the absence of oxygen and therefore can only grow at early stages of fermentation (van Rijswijck et al. 2017).

### Metagenomic deconvolution reveals novel strains and species of bacteria, yeast

In addition to this novel hybrid, we additionally identified and assembled whole genome sequences for three other species of yeast and four species of bacteria. These included the yeasts *Saccharomyces cerevisiae, Dekkera bruxellensis*, and *Pichia kudriavzevii. D. bruxellensis* and *P. kudriavzevii* are both generally associated with spoilage in beer and wine, though *D. bruxellensis* is strongly associated with spontaneous fermentation beers and is prevalent in these cultures after 4 to 8 months of fermentation (Vanoevelen et al. 1977; Verachtert and Iserentant 1995; Spitaels et al. 2014c; Steensels et al. 2015). *D. bruxellensis* is known to have considerable strain variation in chromosome number and ploidy with genome size estimates ranging dramatically; here, the size estimate is comparable to haploid *S. cerevisiae*, considerably smaller than the 20-30 Mb previously estimated (Hellborg and Piskur 2009), although this could partially be explained by incomplete assembly.

The *S. cerevisiae* sequence appears to be closely related to the Canadian beer strain BE018 and the domesticated European wine cluster (Strope et al. 2015; Gallone et al. 2016) (**Figure 3**), though analysis of allele frequencies indicated multiple strains may be present, or that the strain varies in ploidy. Although the original wort inoculation was spontaneous, at various points over the course of the Old Warehouse production the barrel was topped off with other beers which were brewed using commercial ale strains. At this point it is unclear if the *S. cerevisiae* isolated from this sample was initially present, or derived from these later additions.

**Figure 3:**
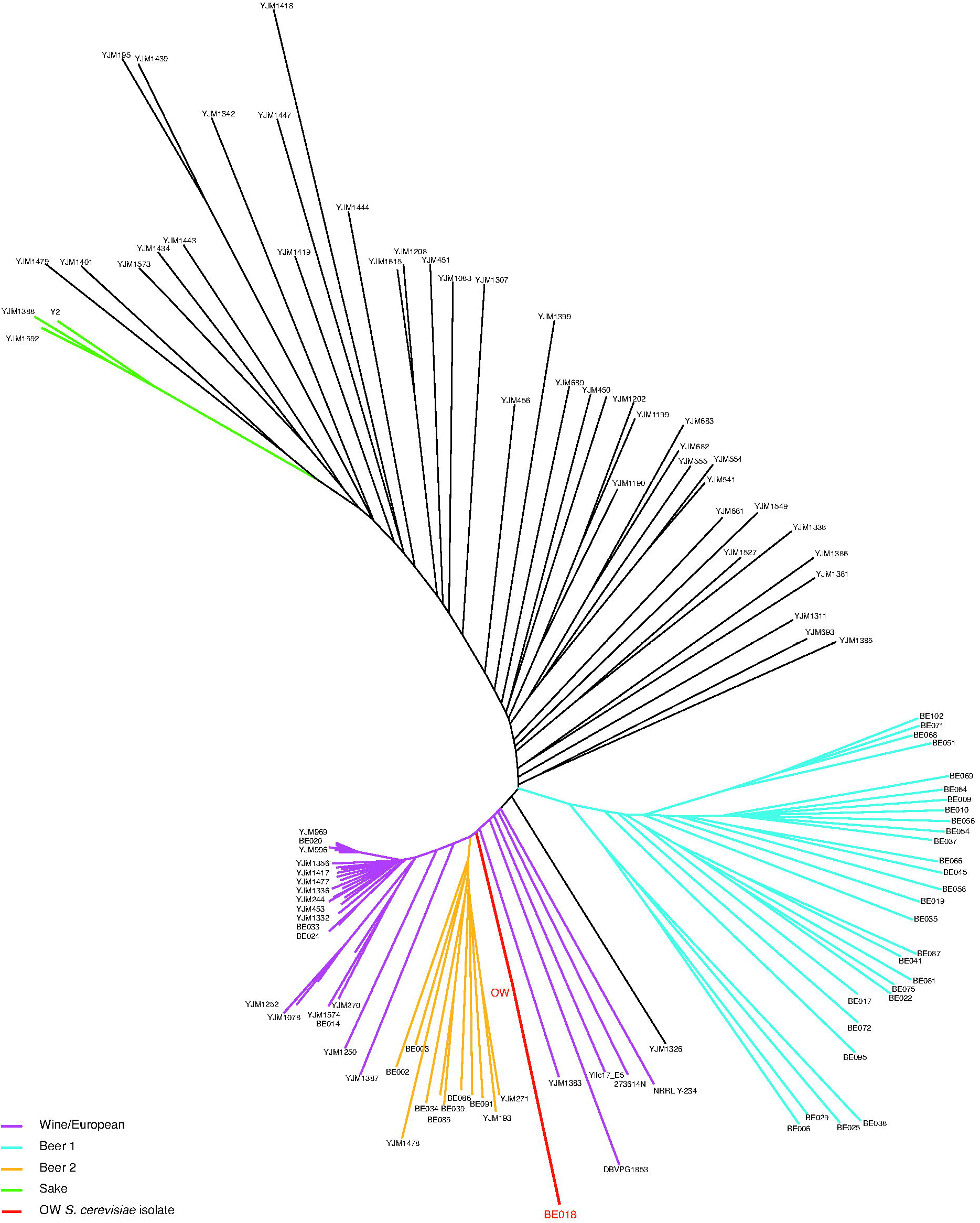
Phylogeny of *S. cerevisiae*. Neighbor-joining tree of *S. cerevisiae* strains representative of the species diversity (Strope et al. 2015; Gallone et al. 2016) and OW (for Old Warehouse) *S. cerevisiae* strain (red).

Consistent with previous studies of spontaneous fermentation beers, we observe acetic acid bacteria (AAB) *Acetobacter pomorum* and a previously unidentified species of Acetobacter (OWB1), and lactic acid bacteria (LAB) *Pediococcus damnosus* and a previously unidentified species of Lactobacillus (OWB2), similar to *L. brevis*. Spitaels *et al*. recently identified two new species of AAB in a traditional Belgian lambic fermentation: *Acetobacter lambici* and *Glunobacter cerevisiae* (Spitaels et al. 2014a; Spitaels et al. 2014b), although we are unable to determine if OWB1 is related to *A. lambici* at this time. Both OWB1 and OWB2 have beer spoilage genes as identified by Fujii *et al*. (Fujii et al. 2005), who identified a locus specific to a wide variety of beer-spoilage strains including *P. damnosus*, *L. collinoides*, *L. coryniformis*, and *L. brevis*.

## Discussion

Here, we leverage Hi-C to characterize a metagenomic sample isolated from a spontaneously inoculated beer from Epic Ales in Seattle, Washington. The method identified the yeasts *D. bruxellensis, P. kudriavzevii, S. cerevisiae*, and a novel *Pichia* hybrid composed of *P. membranifaciens* and an unidentified, related species. We also observed the AAB *Acetobacter pomorum* and a novel Acetobacter species, and the LAB *Pediococcus damnosus* and a novel Lactobacillus species. We present genome assemblies for each, without culturing or any a priori knowledge of community makeup. This represents one of the first studies to deconvolute an unknown mixed sample without culturing (see also (Marbouty et al. 2014; Marbouty et al. 2017)), and demonstrates the ability of this method to both identify novel species, and of particular interest, identify novel hybrids.

### Spontaneous inoculation brewing

The traditional Belgian lambic style of fermentation dates back hundreds of years (Keersmaecker 1996), and has recently gained popularity in the American craft brewery community. Phenotypic identification and culturing, and more recently various genetic methods, have illustrated the microbial progression of these beers, which typically lasts 1-3 years. The first phase lasts for 30-40 days and is dominated by Enterobacteriaceae (Vanoevelen et al. 1977; Martens et al. 1991; Martens et al. 1997; Bokulich et al. 2012; Spitaels et al. 2014c); this is followed by the main fermentation, characterized by *Saccharomyces* yeasts (including *S. cerevisiae, S. pastorianus*, and *S. uvarum)* and lasts 3-4 months. The acidification phase succeeds fermentation and is characterized by the increasing isolation of LAB, and *Dekkera* species after 4 to 8 months of fermentation. Around 10 months, LAB start to decrease as the wort is gradually attenuated. AAB are isolated throughout the brewing process.

In comparison to previous spontaneous inoculation studies, which typically sample over the course of 1-3 years, we only sampled one batch at a single time-point. The sample was obtained after new wort was added to a used wine barrel, from which several prior batches of the “Old Warehouse” beer had already been bottled. Thus, the mixed population we extracted had likely been established for several years in various configurations. We recapitulate some of the previous findings, and also identify several novel strains of bacteria. The diversity of AAB, LAB, and other yeasts found in cultures at various timepoints in other samples can be explained by multiple possibilities. It’s possible that some species are present at undetectable levels in our sample, or that more species are present in the slurry of the barrel, or are embedded in the barrel itself, that were not captured with our sampling method, which only extracted actively growing culture from the aqueous phase. Furthermore, many researchers have found that microbiota are specific to a given environment (Knight et al. 2015; Jara et al. 2016), so differences between different breweries are to be expected. For example, traditional lambics are brewed in breweries in Belgium that produce no other style of beer, whereas in the United States, a different community of yeast and bacterial species are established in breweries that may be experimenting with lambic style fermentation (Bokulich et al. 2012).

While *S. cerevisiae* appears chiefly responsible for fermentation, mixed fermentations with non*-Saccharomyces* yeasts in wine and cider have been shown to produce desirable aroma and flavor complexity (Rojas et al. 2001; Clemente-Jimenez et al. 2005; Viana et al. 2008; Ye et al. 2014; van Rijswijck et al. 2017). The yeast *D. bruxellensis*helps characterize the “funky” flavor of spontaneous inoculation beers; it produces phenol compounds, described as clove-y or medicinal, and smaller compounds often described as barnyard or horse blanket. *D. bruxellensis* and *S. cerevisiae* actually share many important brewing traits, such as the ability to produce high ethanol levels, high tolerance towards ethanol, and the ability to grow without oxygen and in acidic environments (Rozpedowska et al. 2011), but they diverged approximately 200 million years ago, and are thought to have independently evolved these phenotypes (Woolfit et al. 2007). While *S. cerevisiae* outcompetes *D. bruxellensis* when sugar is abundant, the roles are changed at the end of fermentation, when sugar is depleted and only low amounts of nitrogen are available (Nardi et al. 2010).

*D. bruxellensis* is more closely related to the *Pichia* species than it is to the *Saccharomyces* yeasts (Woolfit et al. 2007; Piskur et al. 2012). Prior to reclassification, the *Pichia* genus included more than 100 species, but has been refined to 20, including the type species of the genus *P. membranifaciens*, and the distantly related *P. kudriavzevii* (Kurtzman et al. 2008; Kurtzman 2011). *P. membranifaciens* and *P. kudriavzevii*are most commonly associated with food and beverage spoilage (Kurtzman et al. 2011; Chan et al. 2012), but seem to play a role in various fermentations (Lachance 1995; Daniel et al. 2009; Silva et al. 2009; N'Guessan K et al. 2011; Del Monaco et al. 2014; Spitaels et al. 2015; Stefanini et al. 2016a; Varela and Borneman 2017). A study of wine deacidification showed that *P. kudriavzevii* increased a wine’s fruity aroma and could degrade L-malic acid, increase the pH, and produce low levels of ethanol, important levels of glycerol, and acceptable amounts of acetic acid (Del Monaco et al. 2014). *P. membranifaciens* is repeatedly isolated from lambic and gueuze beers (Spitaels et al. 2014c; Spitaels et al. 2015). Attempts to ferment with the *Pichia* hybrid discovered in this sample suggest that it does not significantly contribute to ethanol production, nor did it change the acidity of the wort; however, it might contribute flavor compounds and/or act in concert with other members of the community. Though recent work suggests sensory profiles of beers fermented with *Pichia* yeasts are undesirable (Osburn et al. 2017), we did not detect an obviously objectionable profile.

### Conclusions

The discovery of a new interspecific hybrid from a brewing environment adds to an increasing number of similarly documented cases, raising several interesting questions. Are there particular conditions in brewing environments that select for hybridization events? For example, a recent study found that chemical-physical conditions in the gut of wasps selected for hybridization events between *S. cerevisiae* and *S. paradoxus* (Stefanini et al. 2016b). Brewing environments could also simply be enriched in both population size and diversity of species, thus presenting more opportunity for hybridization than their natural environments. Perhaps previously established hybrids have a selective advantage in brewing environments, which there is some evidence for with *Dekkera* hybrids in wine environments (Borneman et al. 2014). Alternatively, the explanation could simply be sampling bias: yeast hybrids happen in many places but breweries and wineries are heavily sampled. These hypotheses are not mutually exclusive, and present a rich field for further study. With new detection tools, like the method presented here, we have the ability to identify new hybrids from a variety of taxa and environments, thus providing a critical tool to understanding hybrid biology and ecology.

## Acknowledgements

We thank Benjamin Vernot for initiating the collaboration with Epic Ales. We appreciate the feedback we received from two anonymous reviewers, and from many brewers, particularly Matthew L. Bochman, Ron Extract, and Tom Schmidlin. This work was supported by National Science Foundation grant 1516330 (M.J.D.), National Institute of Health (NIH)/National Human Genome Research Institute (NHGRI) grant T32HG000035 (J.N.B.), NIH/NHGRI grant HG006283 (J.S.), NIH/National Institute of General Medical Sciences grant P41 GM103533 (M.J.D.), DOE-LBL-JGI grant 7074345/DE-AC02-05CH11231 (J.S.), and NIAID grant 1R43AI122654 (I.L.). I.L. was also supported by the UW Commercialization Gap Fund and Commercialization Fellows Program. M.J.D. is a Fellow in the Genetic Networks program at the Canadian Institute for Advanced Research and is supported in part by a Faculty Scholar grant from the Howard Hughes Medical Institute.

**COI:** JNB, IL, J. Shendure, and MJD have financial interests in Phase Genomics, a company that has commercialized the Hi-C genome assembly technique. IL is a current employee of Phase Genomics.

